# Mitochondrial genome variation affects the mutation rate of the nuclear genome in *Drosophila melanogaster*

**DOI:** 10.1101/122234

**Authors:** Jonci N. Wolff, M. Florencia Camus, Damian K. Dowling, Björn Rogell

## Abstract

Mutations are the raw material for evolutionary change. While the mutation rate has been thought constant between individuals, recent research has shown that poor genetic condition can elevate the mutation rate. Mitonuclear genetic conflict is a potential source of poor genetic condition, and considering the high mutation rate of mitochondrial genomes, there should be ample scope for mitochondrial mutations to interfere with genetic condition, with concomitant effects on the nuclear mutation rate. Moreover, because theory suggests mitochondrial genetic effects will often be male-biased, such effects could be more strongly felt in males than females. Here, by mating irradiated male *Drosophila melanogaster* to isogenic females bearing six distinct mitochondrial haplotypes, we tested whether mitochondrial genetic variation affects DNA repair capacity, and whether effects of mutation load on reproductive function are shaped by interactions between sex and mitochondrial haplotype. We found mitochondrial genetic effects on DNA repair, and that the mutational variance of reproductive fitness was higher in males bearing haplotypes characterized by high female fitness. These results suggest that mitochondrial genome variation may affect the mutation rate, and that induced mutations interact more strongly with male than female reproductive function. The potential for haplotype-specific effects on the nuclear mutation rate has broad implications for evolutionary dynamics, such as the accumulation of genetic load, adaptive potential, and the evolution of sexual dimorphism.

## Background

Mutation is the primary source of all genetic variation, both adaptive and deleterious. Generally, *de novo* mutations are detrimental, and high mutation load will thus reduce mean fitness, posing a threat to individual and population viability (Keightley and Lynch 2003). However, mutations are also the source of standing genetic variation, enabling populations to mount an evolutionary adaptive response to sudden environmental change and changing habitats. Mutations thus stand at the core of a population’s ability to adapt and to evolve. Understanding the factors that define the frequency at which new heritable mutations arise - the mutation rate - is thus of fundamental interest to evolutionary biology. Mutation rates are known to vary between species, and while most mechanisms that define the mutation rate are not yet fully resolved (Baer et al. 2007), one key component is the capability of organisms to repair damage incurred to their DNA (Sniegowski et al. 2000).

The prevailing paradigm for the analysis of mutations, from a population genetics perspective, is that the genome-wide mutation rate between individuals in a population is relatively constant (Drake et al. 1998). However, emerging research suggests inter-individual variability and context-dependency of mutation rates a more realistic scenario (Maklakov et al. 2013; Sharp and Agrawal 2016). Early research in the fruit fly *Drosophila melanogaster* (*D. melanogaster*) indicated condition-dependency of individual mutation rate by revealing sensitivity of DNA repair capacity to the thermal (Sollunn and Strömnæs 1964), nutritional (Clark 1972), and genomic environment (Würgler and Maier 1972). More recently, Maklakov et al. (2013) further found that environmental stress, induced via sexual harassment, reduces the yield of dominant lethal mutations in fruit fly. Such a pattern suggested that individuals affected by environmental stressors show increased investment in repair mechanisms, albeit at the cost of reduced reproduction. Also testing condition-dependence of DNA repair, Agrawal et al. (2008) estimated the number of sex-linked recessive lethals (SLRLs) between low- and high-condition adult fruit fly invoked by dietary restriction during larval stages, and found that low-condition females transmitted more SLRLs than high-condition females, increasing mutational load of descendants of low-condition females. Interestingly, under similar experimental conditions it was found that relative adult usage of three DNA repair pathways differed between low- and high-condition females in response to nutritional environments experienced during early developmental stages (Wang and Agrawal 2012). Distinct DNA repair pathways differ in in their mutational consequence, and may thus underpin observed discrepancies in mutation load.

Condition-dependence of mutation rates has also been observed in response to genetic stressors. A mutation accumulation (MA) experiment found that if fruit fly were left to accrue more deleterious mutations, thereby reducing genetic condition, their descendants displayed an accelerated decrease in viability when compared to individuals from a control population, a finding consistent with an elevated mutation rate (Ávila et al. 2006). Similarly, Sharp *et al*. (2012) constructed MA lines of varying genetic fitness, and found that poor-quality genotypes, bearing deleterious alleles (unrelated to effect DNA repair or replication), caused a marked increase in spontaneous mutation rate. Analyzing the mutation spectrum in MA lines further revealed differences in point mutations, small indel rates, and rates of gene conversion when compared to control lines (Schrider et al. 2013; Sharp and Agrawal 2016). The contrasting pattern in mutation rates between genotypes of high and low genetic quality were underpinned by the use of different condition-dependent DNA repair pathways, and were associated with body mass as measure of overall fitness (Wang and Agrawal 2012; Sharp and Agrawal 2016). Intriguingly, rather than using artificial constructs, some of these experiments relied on genetic variation maintained within the same natural population, suggesting that - based on genetic condition - inter-individual mutation rates within populations may be much more variable than previously thought.

Nuclear genetic variation is perhaps the best-studied mechanism by which genetic condition is affected. However, individual genetic condition also depends on the integrity and function of mitochondrial loci and their epistatic interaction with nuclear loci (Rand et al. 2004). Fundamental biological processes hinge on these interactions, and perturbation of the mitonuclear lineage is known to reduce genetic fitness, critically affecting the expression of a broad range of ecologically-relevant life-history traits (Dobler et al. 2014; Wolff et al. 2014; Hill 2015). Standing at the core of cellular function, it is plausible that genetic variation in the mitochondrial genome maintained in populations also affects processes leading to modification of DNA repair capacity - and thus mutation rate - via mitonuclear interactions. Generally, mutation rates are much higher for mitochondrial than nuclear genes, and there should hence be ample scope for mitochondrial *de novo* mutations to interfere with genetic condition via mitonuclear interactions and incompatibilities. It is thus possible that mitochondrial genetic diversity, and ensuing mitonuclear interactions, play a decisive role in the rate at which novel mutations accumulate in the nuclear genome.

Effects of mutations are often sex-specific, and several studies in *Drosophila* have found that selection against novel mutations is stronger in males than in females (Pischedda and Chippindale 2005; Stewart et al. 2005; Sharp and Agrawal 2008). In this context, mutations in the mitochondrial genome are of particular interest since the maternal mode of mitochondrial inheritance limits selection on the mitochondrial genome to females. This genetic asymmetry should in theory lead to the build-up of a male-specific detrimental mutational load (Frank and Hurst 1996; Beekman et al. 2014), often referred to as *Mother’s curse* (Gemmell et al. 2004). Indeed, a growing body of empirical evidence has supported this evolutionary prediction (Smith et al. 2010; Innocenti et al. 2011; Camus et al. 2012; Yee et al. 2013; Camus et al. 2015; Patel et al. 2016), although the generality of a male-bias across metazoans remains open to question (Mossman et al. 2016). Nonetheless, if male genomic integrity is indeed more compromised than that of females, it is possible that perturbation of mitonuclear epistatic interactions via *de novo* mutations in the nuclear genome may be more strongly felt in males than females.

Here, we tested whether low genetic condition induced by mitonuclear incompatibilities affects the mutation rate in the fruit fly *D. melanogaster*. Fruit flies offer an exceptional model to test this question because males virtually lack any post-meiotic activity in DNA repair in sperm (Würgler and Maier 1972; Graf et al. 1979). However, DNA damage in mutagenized sperm can be repaired by the maternal repair system post-fertilization in zygotes (Vogel et al. 1985). Mutagenized sperm can thus be used to subject the female DNA repair system to standardized DNA damage by crossing irradiated males to healthy wild-type females. Subsequent changes in egg-hatch rates can therefore be used as a proxy for female DNA repair capacity of lethal-dominant alleles. We have conducted two experiments. In the first experiment, we probed for potential effects of mitochondrial genetic variation on DNA repair capacity utilizing six strains of fruit fly, where each strain harbored one of six distinct mitochondrial haplotypes, expressed alongside an isogenic nuclear background (*w^1118^*; Bloomington #5905). The fly strains bearing distinct mitochondrial haplotypes were divided into two groups based on their genetic condition as previously determined (Florencia Camus; personal communication). Mitochondrial haplotypes with a hatch rate higher than 72% (range: 72.1%-77%) were categorized as high genetic fitness, whereas mitochondrial haplotypes with hatch rates less than 65% (range 64.1% - 64.7%) were categorized as lo, genetic fitness. To achieve this, we crossed irradiation-mutagenized males of the shared isogenic nuclear background to females of each of the six mitochondrial strains and subsequently determined the egg hatch rate of these F_0_ females. For the second experiment, we used F_1_ offspring reared from the first experiment to examine whether the irradiation-induced nuclear mutation load interfered with reproductive functions in males and females. To do so, we determined egg hatch rate for flies derived from crosses of male or female F1 offspring mated to flies of the opposite sex sourced from our untreated isogenic control population (hence carrying half a genome with novel mutation load, and one healthy half of the genome). The inclusion of haplotypes of both low and high genetic fitness allowed us to probe for effects of mitochondrial haplotype and female condition on DNA repair. Together, these experiments hence enabled us to test (1) whether mitochondrial genomic variation affects DNA repair capacity, (2) whether this effect is mediated via female genetic condition, and (3) whether mitochondrial genetic effects on DNA repair are sex-specific in magnitude.

## Methods

### Fly strains

We used six mitochondrial haplotypes, originally sourced from natural populations in Alstonville Australia [ALS], Barcelona Spain [BAR], Israel [ISR], Madang Papua New Guinea [MAD], Sweden [SWE], and Zimbabwe [ZIM]. These haplotypes are delineated by a maximum of 172 mutations, 38 of which lie within the coding region, and 13 of which cause non-synonymous changes in the amino acid sequence (Clancy 2008; Wolff et al. 2015). None of these mutations (nor the genes encoded in the mtDNA) have known involvement in processes of DNA replication or DNA repair. Each of these haplotypes had been placed alongside an isogenic nuclear background, *w^1118^*, creating distinct “mitochondrial strains” (Clancy 2008), and each mitochondrial strain has been maintained in independent duplicates since their inception in 2007 (hereafter “mitochondrial strain duplicates”). Since then, mitochondrial strain duplicates were propagated as separate entities, with virgin daughters of each mitochondrial strain duplicate back-crossed to males of the *w^1118^* strain for a further 70 generations to continually warrant nuclear genomic isogenicity across all mitochondrial strains. The *w^1118^* strain itself is propagated by one pair of full-siblings to maintain isogenicity within the nuclear background. Using mitochondrial strain duplicates as internal controls, the genetic architecture of this model thus allows us to unambiguously partition true mitochondrial genetic from nuclear genetic effects. Furthermore, all mitochondrial strains are density-controlled during breeding and experimental procedures, maintained on a potato-dextrose-agar medium at constant temperature (25.0 ± 0.1°C), and diurnal cycle (12h:12h light:dark). All maintenance and experimental crosses (including those of the parental and great-parental generation) involved age-controlled flies (4 days of age at time of oviposition).

### Irradiation

Irradiation of focal flies was conducted at the Monash Animal Research Platform (MARP), using a Caesium-137-based Gammacell^®^ 40 Exactor (Best Theratronics Ltd., Ottawa, Canada). To determine the appropriate irradiation dosage for our experiments, we conducted a separate dose-effect experiment on males of the w1118 strain. Briefly, we subjected replicated cohorts of ten *w^1118^* males each to increasing levels of irradiation (0, 6, 12, 18, 24 Gray [Gy]; three biological replicates per dosage), and determined the irradiation dose at which the percentage of unhatched eggs ovipositioned by *w^1118^* females mated to irradiated males reached approximately 50%. The required dosage of irradiation was thus determined to be 20 Gy (Figure 1S).

### Hatch Rate Assay

The flies of all of the focal crosses, outlined below under *Experimental Design*, were allowed to mate in cohorts of 30 flies at a 1:1 sex ratio for 24 hours. Thereafter, flies were transferred for ovipositioning into 70 ml specimen containers (with mechanically-perforated bottoms to enable aeration), where the inner lid diameter – covered with potato-dextrose-agar medium - served as an egg-laying pad (part #S5744SU, Techno Plas Pty Ltd, Adelaide, Australia). After 20 hours of ovipositioning, flies were removed, and the eggs were incubated for a further 28 hours for hatching. After incubation, egg-laying pads were transferred to 4°C to halt development, and the number of hatched and unhatched eggs per container was counted.

## Experimental design

### Effects of the mitochondrial haplotype on DNA repair

In the first experiment - conducted on the F_0_ generation - we tested the effect of the mitochondrial haplotype on the capacity for DNA repair. To achieve this, we established ten replicate populations of fruit fly - each consisting of 15 virgin females of a given mitochondrial strain and 15 irradiated (treatment) or 15 non-irradiated (pcontrol) virgin *w^1118^* males - for each strain duplicate and for each of six mitochondrial strains (= 6 mitochondrial strains × 2 duplicate strains × 2 treatments = 24 population categories × 10 replicate populations = 240 populations in total; Figure S2). Control strains were treated as procedural controls, *i.e*. were transported alongside treated strains to and from the irradiation facility. Males and females of the F_0_ generation were combined for a 24-hour mating period immediately after the irradiation treatment of *w^1118^* males. After mating, flies were transferred for 20 hours onto egg-laying pads for ovipositioning. At the end of ovipositioning, egg-laying pads were divided into two equal halves, such that both halves contained approximately equal numbers of eggs. One half was transferred to 4°C after 28 hours of incubation to halt development for the Hatch Rate Assay. The second half was transferred into a new vial containing potato-dextrose-agar medium to generate the F_1_ generation. Importantly, at no time of the experiment were the focal mitochondrial haplotypes, which were the subject of this study, exposed to irradiation (only males were irradiated and they possessed the ‘*w^1118^*, mitochondrial haplotype, which was not transmitted to the offspring, given the maternal inheritance of the mitochondrial genome), thus warranting that all irradiation-induced mutations transmitted to the developing zygotes, following the matings, were limited to nuclear loci.

### Effects of Interactions between mitochondrial haplotype and irradiation-induced mutation load on reproductive function

In the second experiment - conducted on the F_1_ generation - we tested whether the mutational load induced by irradiation of males in the F_0_ generation, interacted with reproductive function of male and female offspring. To test this, we collected ten virgin males and ten virgin females from each of the 240 F_1_ populations, and combined flies of the same sex for each of the 24 population categories (six mitochondrial strains × two duplicate strains × two treatments × two sexes = 48 pools of 100 flies each). We then randomly drew 75 flies from each pool to establish five replicate vials containing 15 virgin flies each for both sexes (five ‘male’ vials, five ‘female’ vials) and for each of the 24 population categories (Figure S3). To each of the 240 vials we added 15 virgin flies of the opposite sex sourced from the *w*^*1118*^ control population, allowed for a 24-hour mating period, and conducted a Hatch Rate Assay on all 240 populations (Figure S3). We used this data to investigate both the changes in mean reproductive fitness across our experimental units, and changes in reproductive variance across our experimental units.

A potential caveat of our study is that this experimental design was not optimized to analyze variances as 15 focal flies were kept in each ‘male’ and ‘female’ vial. As a consequence, the residual variance was defined as the variance across vials rather than across individuals. While this may present a shortcoming of our experimental design, the presence of, for example, one sterile male would decrease vial variance by 1/15 (assuming the same mating probability across all males per vial). We hence viewed vial variances as an acceptable proxy for variances in reproductive success. If males with poor fertility also perform poorly in sexual competition, vial variance values should be conservative, as other more fertile males would have mated with the females. However, it is possible that results were biased if sterile males (producing sterile sperm) are equally successful in sexual competition and if the same does not apply to female vials. This could be the case if females that carry a detrimental mutational load produce fewer eggs (rather than unviable eggs), as the proportion of eggs that hatch per vial could be inflated by increased fecundity of fertile females, producing similar variances across vials (each vial has similar proportion of hatched versus unhatched eggs, as eggs were only laid by healthy females). If this was true, we would expect fewer eggs in vials with irradiated females. We hence tested for differences in total egg count between the treatments using a generalized linear model with a Poisson distribution allowing for overdispersion of the data, and found no significant effects (treatment*mitochondial haplotype interaction: P = 0.88).

## Statistical analysis

### Analysis of differences in means - Effects of the mitochondrial haplotype on DNA repair

We tested the effect of irradiation on DNA repair in two different models; one assessing whether DNA repair differed between specific haplotypes, and one assessing whether female genetic condition that characterized the different haplotypes (high and low) affected DNA repair. In the first model we analysed the proportion of eggs that failed to hatch in crosses where females were mated to irradiated males as a binomial response (number of hatched eggs, number of eggs that failed to hatch) depending on the treatment (irradiated, control) and mitochondrial haplotype (ALS, BAR, ISR, MAD, SWE, ZIM), as well as their interaction. Mitochondrial duplicate was added to the model as a random effect, explicitly nested under mitochondrial haplotype. In the second model, we assessed hatch rate as a function of condition (high, low), treatment, and their interaction. In this model, mitochondrial haplotype (nested under treatment and condition) and duplicate (explicitly nested under haplotype) were added as random effects. In both models, a significant interaction between mitochondrial haplotype (first model) or condition (second model) with irradiation treatment would imply that fly strains harbouring distinct mitochondrial haplotypes differ in their capacity to repair DNA damage.

### Analysis of differences in means - Effects of Interactions between mitochondrial haplotype and novel mutation load on reproductive function

We continued by analysing the effect of the introduced mutations on reproductive function, defined as the ability to produce viable gametes of both males and females. We started with two initial models. One model in which we added treatment, sex and mitochondrial haplotype, as well as all interactions; and a second model in which we added treatment, sex and condition, as well as all interactions. Mitochondrial duplicate, explicitly nested under mitochondrial haplotype, was added as random effect to all models. For the model examining the effect of female genetic condition on offspring reproductive performance, mitochondrial haplotype was added as random effect, explicitly nested under condition. Given the high number of explanatory variables, both initial models were subsequently reduced by removing marginal terms, and where the best fit was identified using lowest deviance information criterion (DIC) score.

### Analysis of differences of variances - Effects of Interactions between mitochondrial haplotype and novel mutation load on reproductive function

Effects of mutation load are likely to change the mean value of fitness components across different experimental units, and also to increase individual variation reflecting differences in individual genetic fitness. Hence, differences in variances could potentially inform how haplotype- or sex-specific mutation loads accumulate. We thus probed for differences in variances depending on the mitochondrial haplotype and irradiation treatment by assessing whether models (with condition, sex, and treatment as fixed effects and mitochondrial haplotype and duplicate as random effects) improved by estimating variances for random effects separately, leading to (i) treatment-specific variances for each random effect (including the residual component), and (ii) sex-specific variances exclusively across vials derived from irradiated parents. We only fitted sex-specific variances for treated vials considering our explicit hypothesis that variances should be higher in vials containing flies derived from irradiated parents. As both female and male offspring of non-irradiated parents revealed low variation in reproductive success, we continued by limiting analyses to offspring data derived from irradiated parents. Within this dataset we tested whether residual variances of offspring reproductive success differed depending on the genetic condition of the female haplotype. In these models, we included sex and condition, as well as their interaction, as fixed effects and mitochondrial haplotype as random effect (nested under condition and irradiation treatment). We did not include duplicate, as duplicate explained a very minor fraction of the variance. We also restricted models to test for condition effects, as testing for specific residual variances for each sex × haplotype combination would lead to a disproportionally large number of parameters considering the structure of the data.

We conducted all analyses using a Bayesian mixed model using the R package MCMCglmm accounting for the overdispersion present in our binomial data (Hadfield 2010). For all models we discarded the first 20,000 iterations as burn-in, followed by 50,000 iterations, sampled every 500 iterations, yielding a posterior sample of 1,000 iterations per chain. Three parallel chains were run for each model and their convergence was assessed using the Gelman-Rubin statistic. All autocorrelations were under an absolute value of 0.11. Flat priors were used on fixed effects and locally uninformative priors were used on random effects.

## Results

### Effects of the mitochondrial haplotype on DNA repair

The proportion of eggs that failed to hatch was affected by both mitochondrial haplotype and treatment (irradiated; control), and their interaction (Table 1). Our results indicate that haplotypes ISR, MAD and ZIM had lower hatch rates than haplotypes ALS, BAR, and SWE, and that hatch rate was negatively affected by irradiation treatment (Figure 1; Table 1). While the general effect of irradiation on hatch rate was stronger than that of the interaction between haplotype and irradiation treatment, our experiments revealed that haplotypes BAR, MAD and ZIM were less affected by irradiation treatment than haplotypes ALS, ISR and SWE. Importantly, haplotypes with both lower and higher baseline fecundity were among haplotypes that were more strongly affected by the irradiation treatment, refuting our hypothesis that observed mitochondrial genetic effects on hatch rate are mediated via impaired DNA repair of low genetic condition females. Indeed, in our analysis of condition effects on DNA repair, the interaction between treatment and condition was not significant (Table 1).

**Figure 1.**
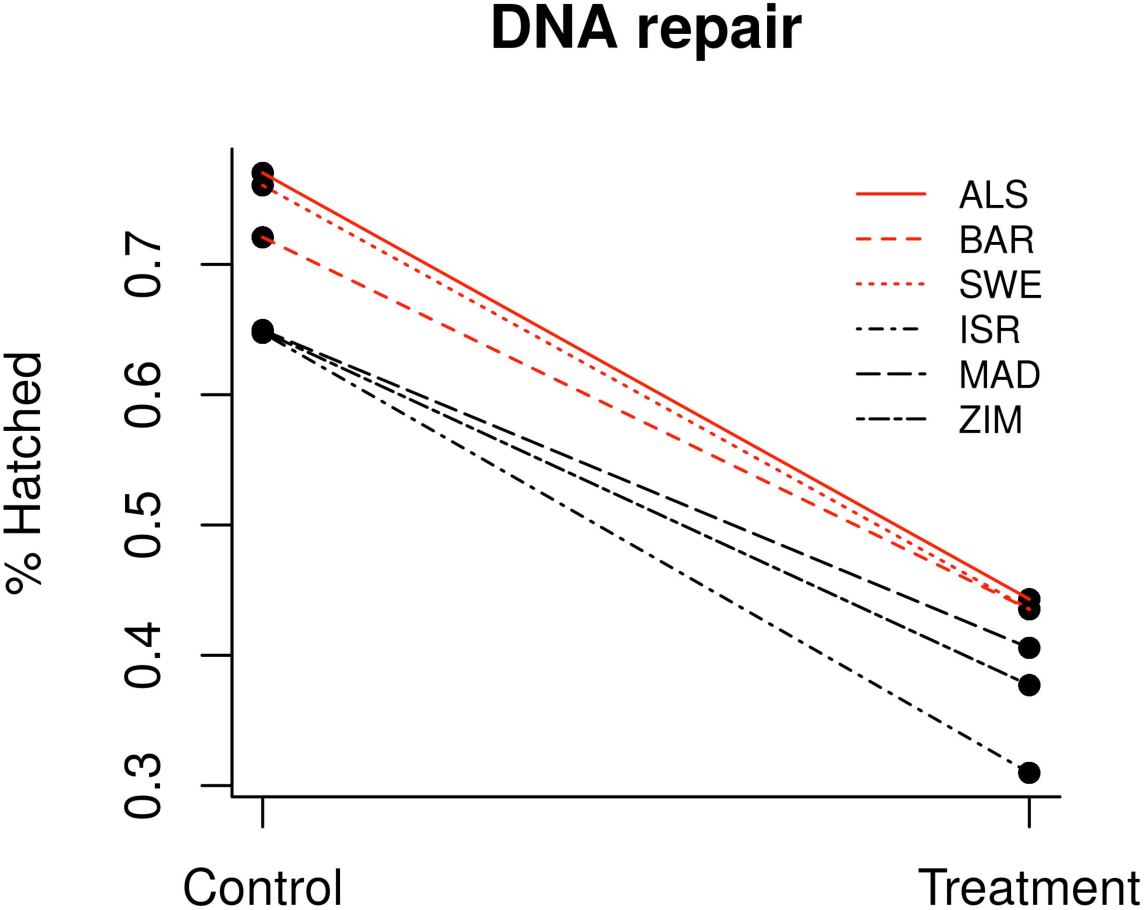
The effect of mitochondrial haplotype and irradiation treatment on F_0_ hatch rate. Hatch rate was determined across six isogenic fly strains bearing six distinct mitochondrial haplotypes (ALS, BAR, ISR, MAD, SWE, ZIM), and where females of the isogenic strains were mated to irradiation-mutagenized males of the isogenic *w^1118^* control strain (Fig. S2). Differences in DNA repair manifest as interactions between haplotype and irradiation treatment. Black lines denote Low condition haplotypes and red lines denote High condition haplotypes.

**Table 1.**
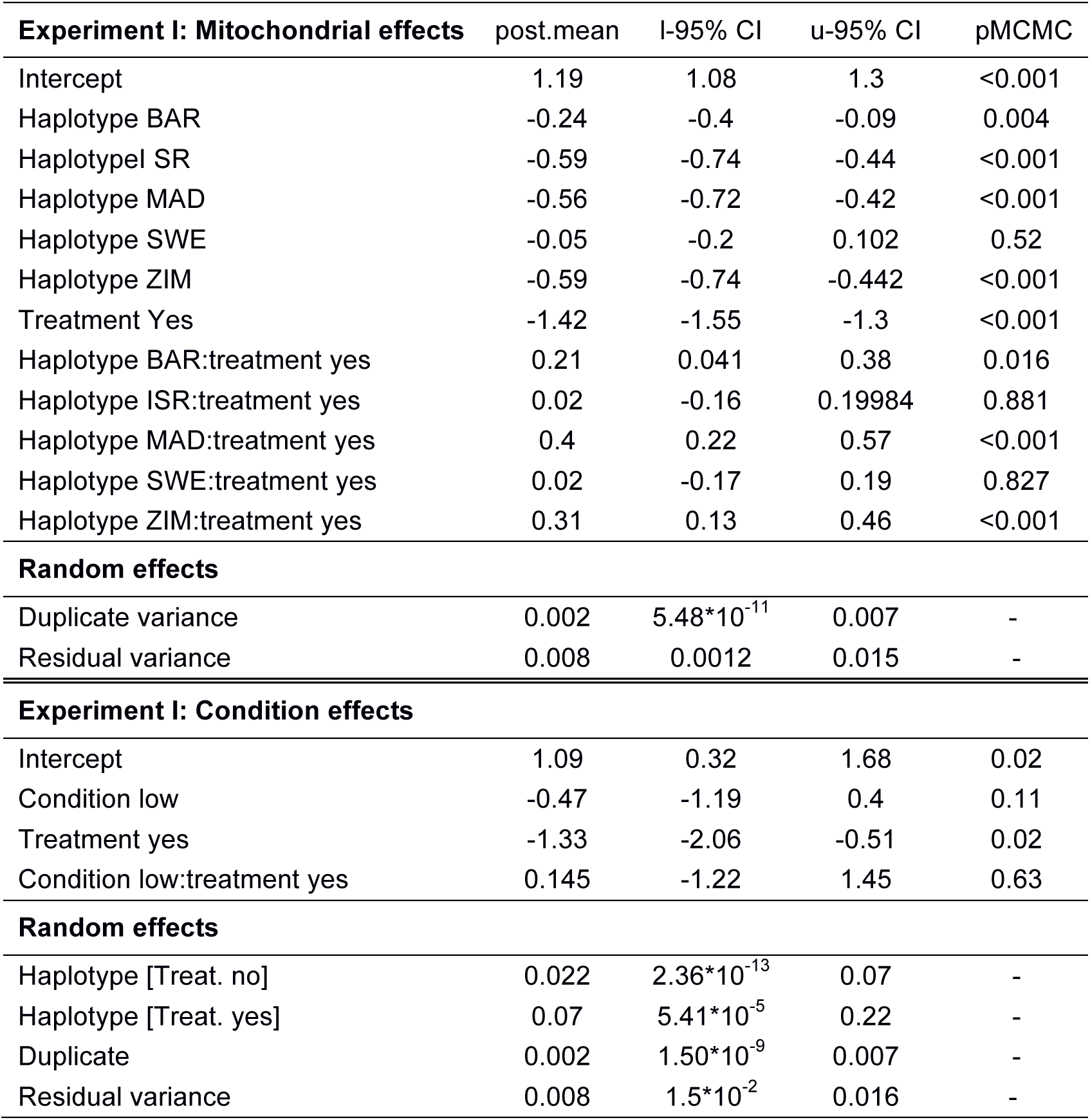
Test of F_0_ hatch rate across six strains of D. melanogaster harboring distinct mitochondrial haplotypes (ALS, BAR, ISR, MAD, SWE, ZIM) mated to irradiation-mutagenized isogenic males. Mitochondrial effects: Results based on estimates using contrast treatments, were the intercept term was set to treatment:no and mitochondrial haplotype ALS. All parameters are contrasted against the intercept. The results indicate that mitochondrial haplotypes BAR, MAD and ZIM are less affected by mating with irradiated males than haplotypes ALS, ISR and SWE. Condition effects:The intercept term was set to *treatment:no* and *condition:high*.

### Effects of Interactions between mitochondrial haplotype and novel mutation load on reproductive function

In the second generation, we found that while both treatment and mitochondrial haplotype affected hatching probability, sex did not have any significant effects on hatching success (Table 2, Figure 2). Neither did we find evidence for interactions between mitochondrial haplotype and irradiation treatment, prompting the removal of all interactions from models with a delta DIC > 3. When analyzing the effect of condition on reproductive success, we found that while both condition and irradiation treatment had significant effects on reproductive success, their interactions were non-significant (Table 2).

**Table 2.** Test of F_1_ hatch rate across six strains of D. *melanogaster* harboring distinct mitochondrial haplotypes (ALS, BAR, ISR, MAD, SWE, ZIM) mated to irradiation-mutagenized isogenic males in F_0_. Mitochondrial effects: Results based on estimates using contrast treatments, were the intercept term was set to *treatment:no* and mitochondrial haplotype ALS. Condition effects: The intercept term was set to *treatment:no* and *condition:high*. All parameters are contrasted against the intercept. Analyses indicate significant effects of mitochondrial haplotype and F_0_ irradiation treatment on hatch rate, but that these effects are not involved in any interactions with the sex or condition of focal flies.

| **Experiment 2: Mitochondrial effects** | post.mean | l-95% CI | u-95% CI | pMCMC |
| --- | --- | --- | --- | --- |
| Intercept | 1.25 | 1.14 | 1.38 | <0.001 |
| Treatment yes | -0.5 | -0.58 | -0.42 | <0.001 |
| Haplotype BAR | -0.13 | -0.28 | 0.058 | 0.13 |
| Haplotype ISR | -0.64 | -0.81 | -0.47 | <0.001 |
| Haplotype MAD | -0.39 | -0.55 | -0.23 | <0.001 |
| Haplotype SWE | -0.11 | -0.28 | 0.065 | 0.2 |
| Haplotype ZIM | -0.31 | -0.49 | -0.15 | 0.006 |
| **Random effects** |  |  |  |  |
| Duplicate variance | 0.0025 | 2.22*10<sup>-8</sup> | 0.0084 |  |
| Residual variance | 0.058 | 0.04 | 0.077 |  |
| **Experiment 2: Condition effects** | post.mean | l-95% CI | u-95% CI | pMCMC |
| Intercept | 1.14 | 0.99 | 1.28 | <0.001 |
| Condition low | -0.36 | -0.49 | -0.22 | <0.001 |
| Treatment yes | -0.47 | -0.62 | -0.3 | <0.001 |
| Sex M | 0.02 | -0.05 | 0.1 | 0.63 |
| **Random effects** |  |  |  |  |
| Haplotype [Treat. yes:Sex F] | 0.009 | 5.43*10<sup>-9</sup> | 0.036 | - |
| Haplotype [Treat. yes:Sex m] | 0.044 | 4.38*10<sup>-9</sup> | 0.16 | - |
| Haplotype [Treatment no] | 0.024 | 6.29*10<sup>-7</sup> | 0.07 | - |
| Duplicate[Treat. yes:Sex F] | 0.0036 | 8.44*10<sup>-10</sup> | 0.014 | - |
| Duplicate[Treat. yes:Sex M] | 0.015 | 7.01*10<sup>-11</sup> | 0.059 | - |
| Duplicate [Treat. no] | 0.0023 | 1.83*10<sup>-9</sup> | 0.01 | - |
| Residual [Treat. no: Sex F: Cond High] | 0.006 | 2.16*10<sup>-4</sup> | 0.018 | - |
| Residual [Treat. y: Sex F: Cond High] | 0.014 | 1.87*10<sup>-4</sup> | 0.0384 | - |
| Residual [Treat. no: Sex M: Cond High] | 0.007 | 2.22*10<sup>-4</sup> | 0.024 | - |
| Residual [Treat. y: Sex M: Cond High] | 0.37 | 0.17 | 0.62 | - |
| Residual [Treat. no: Sex F: Cond low] | 0.0034 | 2.22*10<sup>-4</sup> | 0.01 | - |
| Residual [Treat. y: Sex F: Cond low] | 0.084 | 0.026 | 0.15 | - |
| Residual [Treat. no: Sex M: Cond low] | 0.0033 | 1.53*10<sup>-4</sup> | 0.01 | - |
| Residual [Treat. y: Sex M: Cond low] | 0.16 | 0.054 | 0.29 | - |

**Figure 2.**
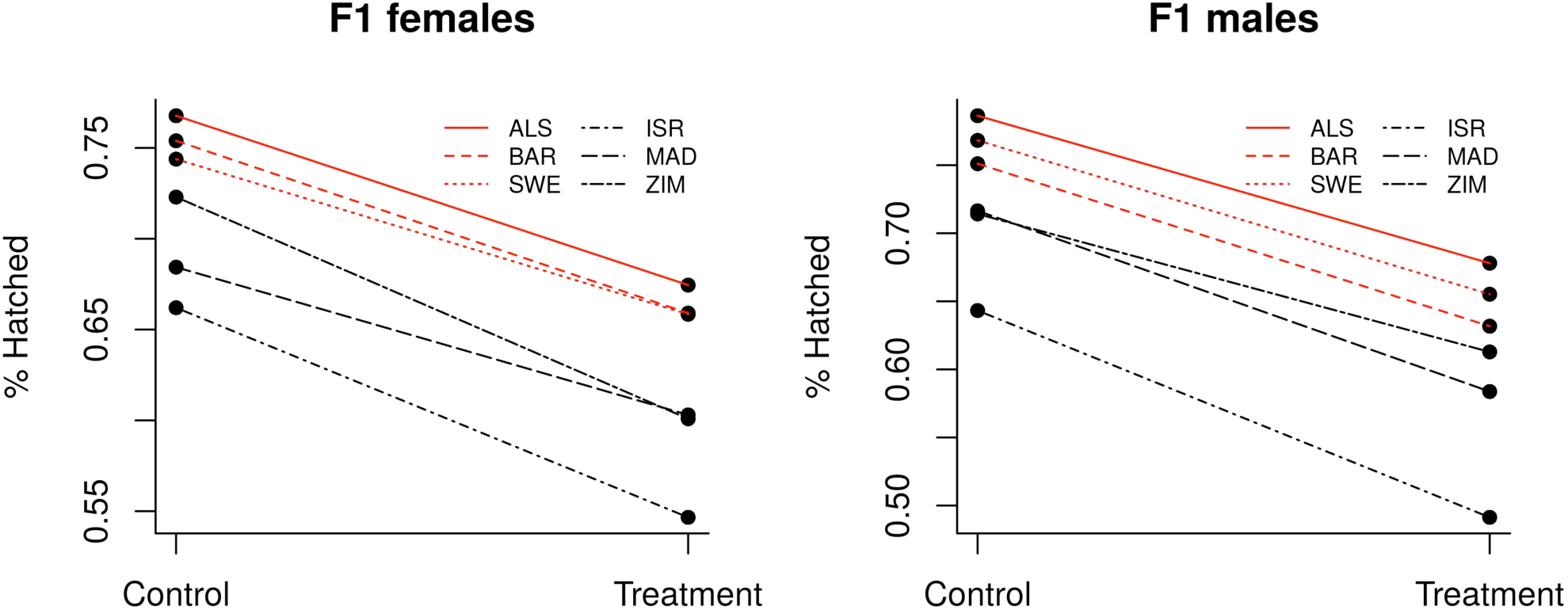
The effect of mitochondrial haplotype and irradiation treatment on F_1_ hatch rate. Hatch rate was determined for groups of mutagenized male or female F_1_ offspring bearing six distinct mitochondrial haplotypes (ALS, BAR, ISR, MAD, SWE, ZIM) and that were mated to untreated flies of the opposite sex sourced from the isogenic *w^1118^* control strain (Fig. S3). Black lines denote Low condition haplotypes and red lines denote High condition haplotypes.

Analyzing variances, we found strong support for the inclusion of different residual variances for flies derived from irradiated flies (Delta DIC = -87), with offspring of irradiated parents exhibiting higher variances. Estimating different variances for irradiation treatment did not improve the fit for mitochondrial variances (delta DIC = 4), or duplicate variances (delta DIC = 1). However, separation into sex- and treatment-specific residual variances did improve the fit (delta DIC = -8), with males derived from irradiated parents exhibiting higher residual variances (Figure 3). When restricting the analysis to irradiated flies only, we found that irradiated males from lines that were characterized by high genetic condition had significantly higher variances than females (delta DIC = 2.8; Figure 3).

**Figure 3.**
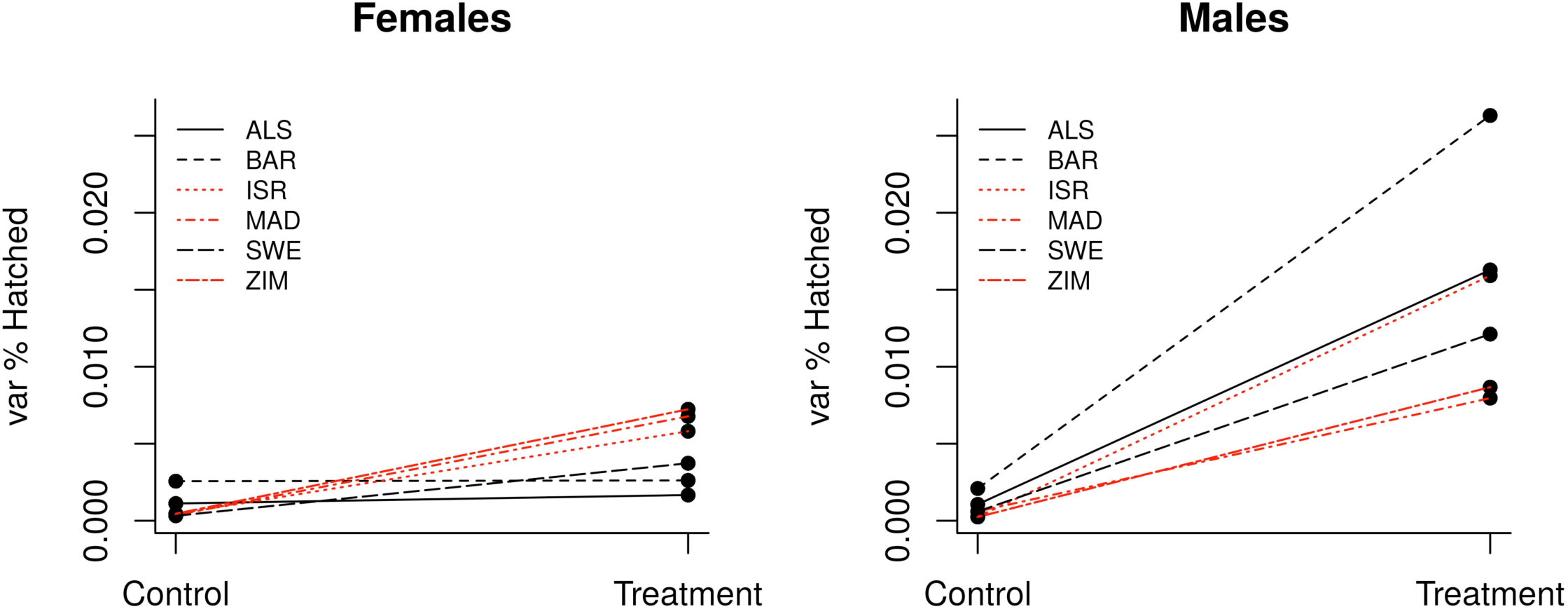
The effect of mitochondrial haplotype and irradiation treatment on variation in F1 hatch rate. Variation in hatch rate for groups of mutagenized female or male F_1_ offspring bearing six distinct mitochondrial haplotypes (ALS, BAR, ISR, MAD, SWE, ZIM) mated to untreated flies of the opposite sex sourced from the isogenic *w^1118^* control strain (Fig. S3). Black lines denote Low condition haplotypes and red lines denote High condition haplotypes.

## Discussion

Mitochondrial genome variation is known to affect genetic condition - and thus organismal fitness - via mitonuclear interactions and genetic conflict within mitonuclear gene complexes (Burton et al. 2006; Ellison and Burton 2006; Dowling et al. 2007a; Dowling et al. 2007b; Barreto and Burton 2013; Dobler et al. 2014; Wolff et al. 2014). Because the nuclear mutation rate has been found to be affected by poor condition invoked by environmental and genetic stressors (Agrawal and Wang 2008; Sharp and Agrawal 2012; Wang and Agrawal 2012; Maklakov et al. 2013), it appears likely that condition-effects induced by mitonuclear incompatibilities might therefore play a significant role in defining the rate at which mutations accumulate in the nuclear genome. Indeed, subjecting females to a standardized mutation load through mutagenized sperm in our first experiment, we show that female capacity to repair *de novo* mutations was contingent on the mitochondrial haplotype. Mitochondria play a crucial role in cell metabolism and cellular respiration (Shadel and Horvath 2015), and perturbation of mitochondrial function via mitonuclear genetic conflict is known to affect physiological state (Arnqvist et al. 2010; Barreto and Burton 2013; Latorre-Pellicer et al. 2016; Wolff et al. 2016a). Our results thus indicate that changes in physiological state invoked by mitonuclear conflict may extend to changes in the susceptibility and the capacity to repair novel mutations. Contrary to our expectation, however, comparing hatch rate between females of high versus low genetic condition after irradiation treatment, we found that observed mitochondrial genetic effects on DNA repair were not condition-dependent on female reproductive fitness. This finding indicates that, while both female fertility (*e.g*., condition) and DNA repair capacity are contingent on the mitochondrial haplotype, the mitonuclear epistatic interactions modulating DNA repair capacity and female fertility differ between these traits, and that the mitochondrial genetic effects on DNA repair are not linked to effects on female condition.

Our finding that DNA repair capacity is modulated by the mitochondrial haplotype provides valuable insight into the mechanisms that shape the mutation rate, and that subsequently feed into processes of adaptation and ultimately speciation. A role for mitochondrial genome variation in modulating nuclear mutation rate is particularly interesting considering the high genetic diversity and mutation rate heterogeneity generally observed for mitochondrial genomes (Brown et al. 1979; Denver et al. 2000; Bazin et al. 2006; Nabholz et al. 2008; Nabholz et al. 2009). Direct measurement of mutation rate across 200 generations of spontaneous mutation accumulation in *D. melanogaster*, for example, estimated the overall mitochondrial mutation rate to be between 10 to 70 times higher than the nuclear mutation rate depending on the nature and strand identity of mutations (Haag-Liautard *et al*. 2008). Major mechanisms thought to underpin the high mutation rate and genetic diversity of mitochondrial genomes are the mutational damage incurred via copying errors during the replication process of mitochondrial genomes (Larsson 2010; Ameur et al. 2011; Kauppila and Stewart 2015), and to a lesser extent, the mutational damage incurred via the leakage of reactive oxygen species (ROS) generated by processes of cellular respiration that take place in mitochondria (Cooke et al. 2003; Balaban et al. 2005; Itsara et al. 2014). Our finding that mitochondrial genetic variation can affect nuclear mutation rates thus suggests that the mechanisms that underpin high mitochondrial mutation rates may ultimately also be drivers of nuclear genomic diversity, if mitochondrial haplotypes interact with DNA repair capacity. Considering the high mtDNA diversity in natural populations (Maklakov et al. 2006; Dowling et al. 2007b), our results further suggests that mitochondrial genetic effects on nuclear mutation rates should be observable between individuals of the same population, and that mutation rate heterogeneity within populations may be much more common and pronounced than previously thought. Support for this notion comes also from recent research that found condition-dependence of individual mutation rate in fruit fly in response to genetic or environmental stressors, such as food scarcity or sexual harassment, impacting on individual physiological state, and consequently mutation rates (Agrawal and Wang 2008; Sharp and Agrawal 2012; Wang and Agrawal 2012; Maklakov et al. 2013).

In our second experiment we tested whether reproductive fitness of offspring bearing novel mutation load, attributable to an inherited copy of an irradiated haploid genome, differed between the sexes, and whether sex-specific variation in reproductive success was higher in low than high condition females. While sex and mitochondrial haplotype did not have any significant effects on mean reproductive success, this test revealed that male offspring derived from high condition females exhibited higher residual variances in hatch rate than their female siblings. This may suggest that specific mutations interacted with the mitochondrial haplotype in a sex-specific manner, and that these effects were contingent on the condition of female mitochondrial haplotype. Likely, individual variation in mutation effects on reproductive function exhibited by each male differed between high and low genetic condition strains, suggesting that males derived from high condition females harbored nuclear mutations that interacted more strongly with mitochondrial loci than the mutations harbored by males derived from low condition females. While this finding was unexpected, the key to understanding this pattern is likely to lie within the uniparental mode of mitochondrial inheritance. In this regard, our observations are to some extent congruent with patterns of male-biased mitochondrial genetic effects as suggested by the *Mother’s curse* hypothesis (Frank and Hurst 1996; Gemmell et al. 2004). Under this hypothesis, mitochondrial traits are not under direct selection in males, enabling the accumulation of mutations in mitochondrial genes that are either male-harming or sexually antagonistic in effect (male-harming, female-beneficial), and which are known to negatively affect male reproductive traits (Clancy et al. 2011; Yee et al. 2013; Dowling et al. 2015; Patel et al. 2016; Wolff et al. 2016b). It is thus possible that, if the mitochondrial haplotypes characterized by high female condition harbor a sexually antagonistic mutation load, that effects of genetic challenge via irradiation-induced mutation load may be more strongly felt in strains of high than low condition haplotypes. Alternatively, it is also conceivable that condition-dependent male variances were driven by differences in the strength of female germ line selection. However, effects of germ line selection would have to be both strong, and differ between haplotypes to explain observed differences between males derived from high and low condition females.

Another potential mechanism to explain observed sex-specific differences in variances are interactions between mitochondrial haplotype and Y chromosome. Contrary to mitochondria, the Y chromosome is inherited strictly paternally, and is thus subject to male-specific selection (Bachtrog 2013). In *D. melanogaster*, the Y chromosome codes for less than 20 male-specific genes, and male fitness has been found strongly affected by Y chromosome polymorphisms in this species (Chippindale and Rice 2001; dos Santos et al. 2015). Moreover, because of its male-specificity in inheritance and gene content, the Y chromosome has been suggested as candidate region to harbour loci that host nuclear modifier alleles that compensate male-harming mutations that may have accumulated under influences of the sex-biased selective sieve described by *Mother’s curse* in mitochondrial genes (Rogell et al. 2014; Dean et al. 2015; Yee et al. 2015). Following our breeding scheme, male offspring carried a Y chromosome harboring a novel mutation load from irradiation-treated male parents, while the X chromosome and one of each of the autosomes were received from an untreated female. Female offspring instead harbored one haploid genome each from an untreated and treated parent, leaving females with a genetically healthy copy for each chromosome. It is thus conceivable that irradiation-induced mutation load in male-specific modifier alleles located on the Y chromosome may underpin increased variances in male reproductive success across different haplotypes.

Together, our findings suggest that mitochondrial genome variation and ensuing mitonuclear interactions and/or incompatibilities may routinely play a pivotal role in the rate at which *de novo* mutations accumulate in the nuclear genome. The implications of mitochondrial haplotype-specific effects on the nuclear mutation rate are broad. Mutations generally decrease fitness, and increases in the mutation rate are thus likely to reduce mean fitness, posing a threat to population viability. Hybrid zones, for example, may be particularly vulnerable to adverse mitonuclear effects on mutation rates. Here, the creation of novel mitonuclear allelic combinations between distantly related individuals are likely to lead to genetic incompatibilities, potentially accelerating the mutation rate (Liu et al. 2017). Conversely, however, populations with high genetic diversity in mitonuclear gene complexes may exhibit increased mutational variation, leading to increased standing genetic variation and therefore increased capacity to adapt and survive in changing environments. This could in theory aid populations in secondary contact zones, where populations of distinct mitochondrial haplotypes could gain increases in nuclear genetic diversity not only by population admixture but also from increased mutation rate. Interestingly, within-population mitonuclear genetic variation also implies that some of a population’s individuals may be disproportionately burdened with mutation load. Individuals with particularly high mutation rate (mutator individuals) could be exceptionally important in times of strong selective pressure and in cases where an adaptive response requires several mutations at once. The transient presence of mutator individuals may thus supplement a population’s potential to adapt, while loading the mutational burden on only few of a population’s individuals, could reduce long-term mutational load and aid escaping the risk of population extinction that would otherwise eventually ensue via mutational meltdown (Agrawal 2002; Shaw and Baer 2011).

In summary, we have provided compelling empirical support that the mitochondrial haplotype affects DNA repair, and therefore mutation rate. The ubiquity of mitochondrial genetic effects on life-history expression further suggests these effects to be pervasive and sizable in magnitude between and within populations (Dowling et al. 2008; Dobler et al. 2014; Alexander et al. 2017). Our data thus provide important new insight into the factors that underpin and determine the mutation rate, and therefore the evolutionary processes that are core to evolutionary diversification and population viability.

## Data accessibility

All raw data are available through the Dryad Digital Repository (www.datadryad.org) #xxx.

## Authors’ Contributions

J.N.W. and B.R. conceived, designed, and analysed the experiments; J.N.W and M.F.C performed the experiment, J.N.W., D.K.D. and B.R wrote the paper.

## Acknowledgements

We thank Belinda Van Leeuwen and Monique Centrone at the Monash Animal Research Platform (MARP) for their help to conduct irradiation experiments. We also thank David Clancy for generously providing the fly strains in 2007.

## Supplementary Material

**Figure S1:**
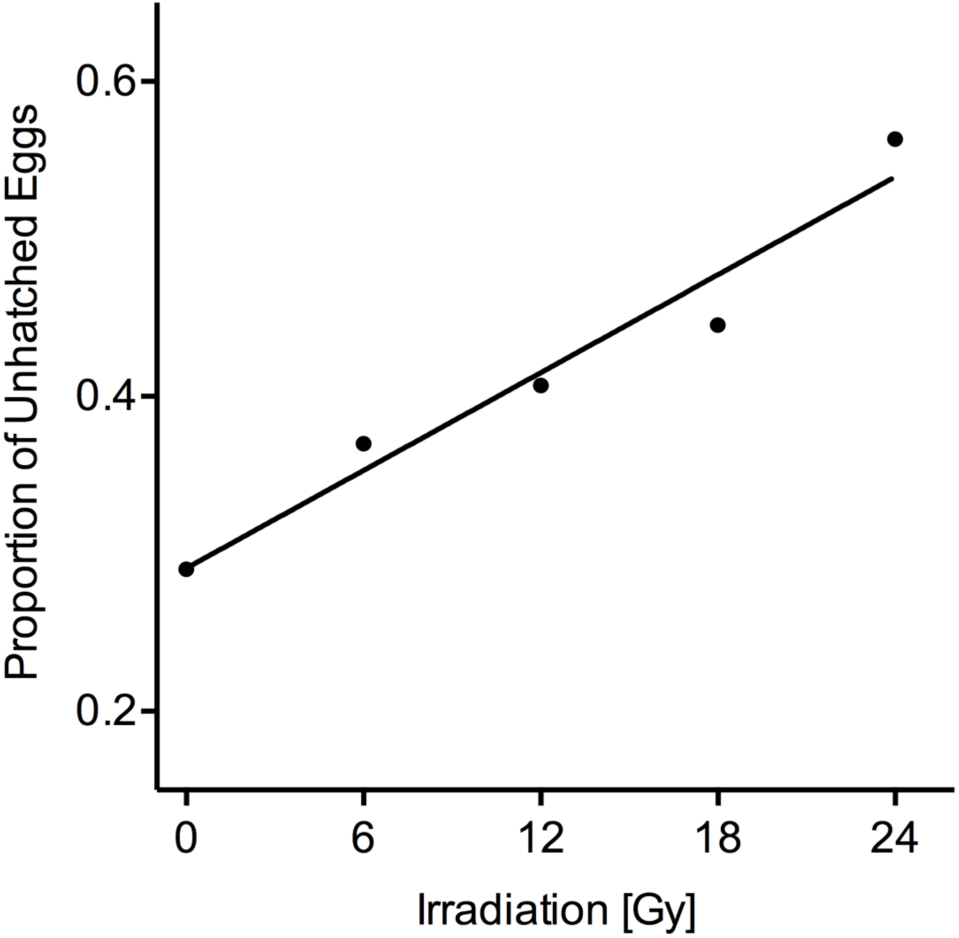
The effect of irradiation on the proportion of unhatched eggs of *Drosophila melanogaster w^1118^* females mated to *w^1118^* males exposed to increasing irradiation dosage.

**Figure S2:**
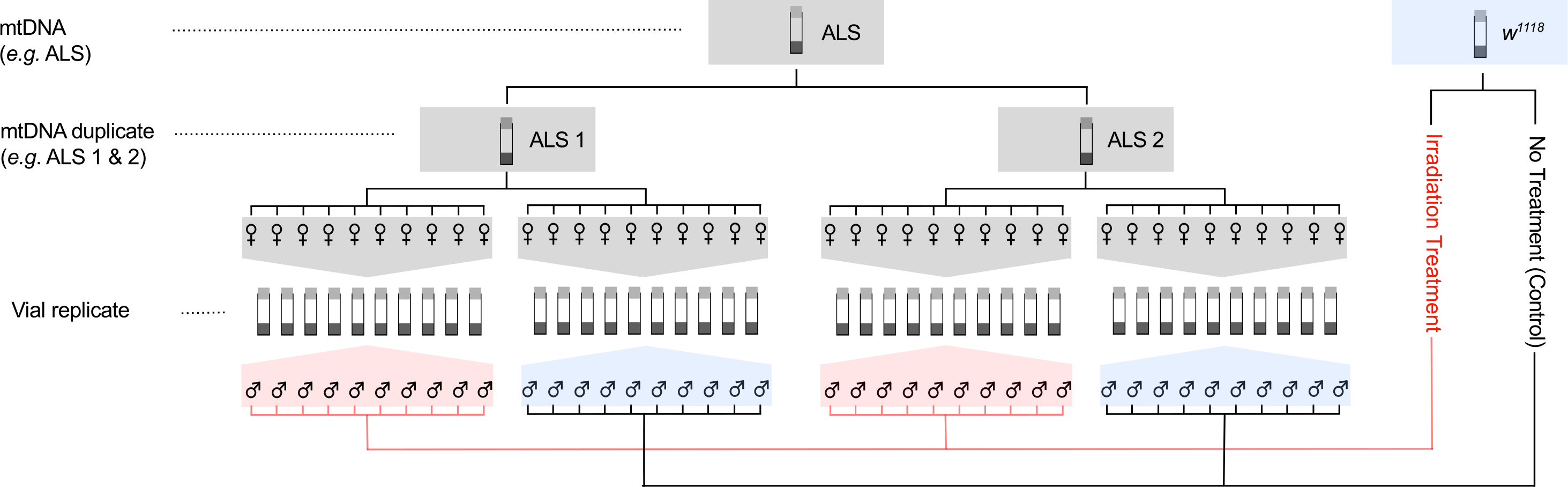
Experimental breeding scheme for the F_0_ generation. Mitochondrial strains (Alstonville [ALS], Barcelona [BAR], Israel [ISR], Madang [MAD], Sweden [SWE], Zimbabwe [ZIM]) were maintained in independent duplicates since 2007 [*e.g*. ALS 1 & 2). For each line duplicate, we collected 300 virgin females for 10 replicates with 15 virgin females each for both the control and treatment cohort. To each of the replicates containing 15 virgin females, we added 15 *w^1118^* virgin males (irradiated for treatment group, non-irradiated for control group), and allowed flies to mate for 24 hours before commencement of F_0_-ovipositioning

**Figure S3:**
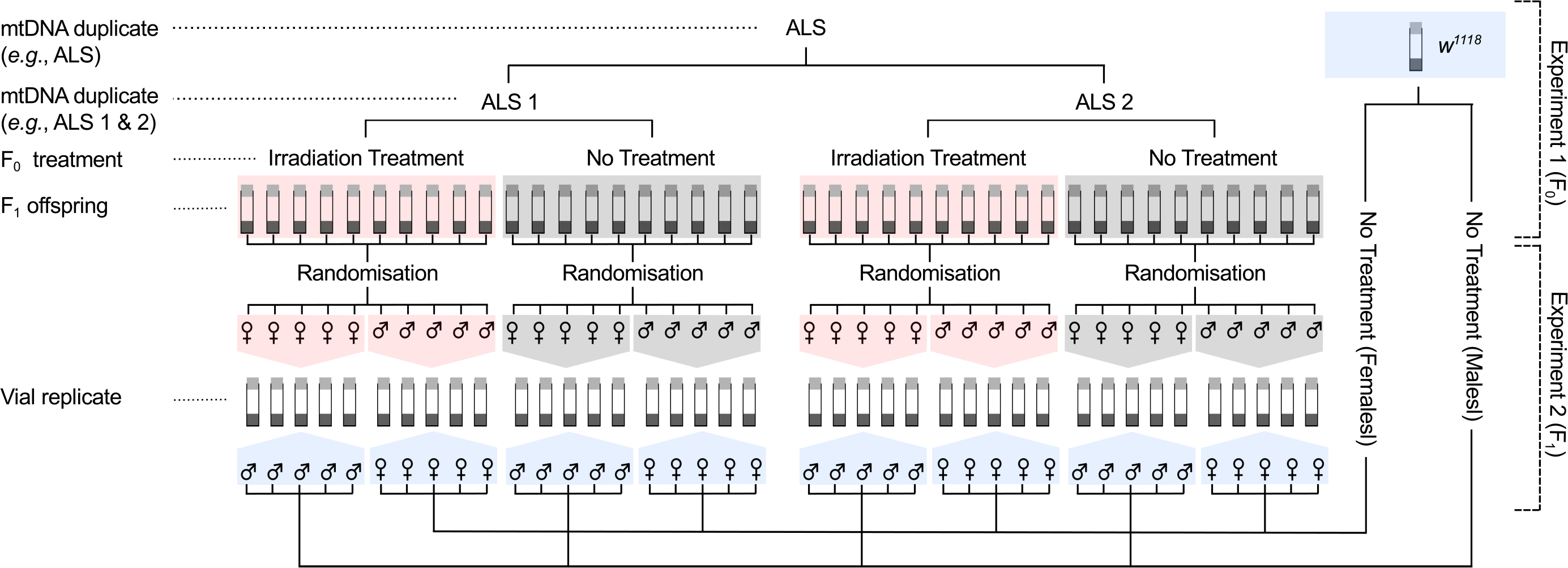
Experimental breeding scheme for the F1 generation. 50% of eggs ovipositioned by the F0 generation (see also Figure S1) were left to develop until eclosion for both duplicates of each of six mitochondrial strains (Alstonville [ALS], Barcelona [BAR], Israel [ISR], Madang [MAD], Sweden [SWE], Zimbabwe [ZIM]). For each line duplicate, we collected 150 virgin females and 150 virgin males, and after randomization of these flies within each cohort and sex, we established 5 replicates for each sex containing 15 virgin flies for both the control and treatment cohort. Each vial was then supplemented with 15 virgin flies of the opposite sex sourced from the isogenic *w^1118^* strain. All populations were allowed to mate for 24 hours before commencement of F_1_-ovipositioning.

## References

Agrawal, A. F. 2002. Genetic loads under fitness-dependent mutation rates. J. Evol. Biol. 15:1004–1010.

Agrawal, A. F. and A. D. Wang. 2008. Increased transmission of mutations by low-condition females: Evidence for condition-dependent DNA repair. PLoS Biol. 6:e30.

Alexander, H. K., S. I. Mayer, and S. Bonhoeffer. 2017. Population Heterogeneity in Mutation Rate Increases the Frequency of Higher-Order Mutants and Reduces Long-Term Mutational Load. Mol. Biol. Evol. 34:419–436.

Ameur, A., J. B. Stewart, C. Freyer, E. Hagstrom, M. Ingman, N. G. Larsson, and U. Gyllensten. 2011. Ultra-deep sequencing of mouse mitochondrial DNA: mutational patterns and their origins. PLoS Genet. 7:e1002028.

Arnqvist, G., D. K. Dowling, P. Eady, L. Gay, T. Tregenza, M. Tuda, and D. J. Hosken. 2010. Genetic architecture of metabolic rate: Environment specific epistasis between mitochondrial and nuclear genes in an insect. Evolution 64:3354–3363.

Ávila, V., D. Chavarrías, E. Sánchez, A. Manrique, C. López-Fanjul, and A. García-Dorado. 2006. Increase of the spontaneous mutation rate in a long-term experiment with *Drosophila melanogaster*. Genetics 173:267–277.

Bachtrog, D. 2013. Y-chromosome evolution: emerging insights into processes of Y-chromosome degeneration. Nat. Rev. Genet. 14:113–124.

Baer, C. F., M. M. Miyamoto, and D. R. Denver. 2007. Mutation rate variation in multicellular eukaryotes: causes and consequences. Nat. Rev. Genet. 8:619–631.

Balaban, R. S., S. Nemoto, and T. Finkel. 2005. Mitochondria, oxidants, and aging. Cell 120:483–495.

Barreto, F. S. and R. S. Burton. 2013. Elevated oxidative damage is correlated with reduced fitness in interpopulation hybrids of a marine copepod. Proc. R. Soc. Biol. Sci. Ser. B 280:20131521

Bazin, E., S. Glémin, and N. Galtier. 2006. Population size does not influence mitochondrial genetic diversity in animals. Science 312:570–572.

Beekman, M., D. K. Dowling, and D. K. Aanen. 2014. The costs of being male: Are there sex-specific effects of uniparental mitochondrial inheritance? Philos. Trans. R. Soc. Lond. B Biol. Sci. 369.

Brown, W. M., M. George, Jr., and A. C. Wilson. 1979. Rapid evolution of animal mitochondrial DNA. Proc. Natl. Acad. Sci. U. S. A. 76:1967–1971.

Burton, R. S., C. K. Ellison, and J. S. Harrison. 2006. The sorry state of F2 hybrids: Consequences of rapid mitochondrial DNA evolution in allopatric populations. The American Naturalist 168:Suppl 6: S14–24.

Camus, M. F., D. J. Clancy, and D. K. Dowling. 2012. Mitochondria, maternal inheritance, and male aging. Curr. Biol. 22:1–5.

Camus, M. F., J. B. Wolf, E. H. Morrow, and D. K. Dowling. 2015. Single nucleotides in the mtdna sequence modify mitochondrial molecular function and are associated with sex-specific effects on fertility and aging. Curr. Biol. 25:2717–2722.

Chippindale, A. K. and W. R. Rice. 2001. Y chromosome polymorphism is a strong determinant of male fitness in Drosophila melanogaster. Proc. Natl. Acad. Sci. U. S. A. 98:5677–5682.

Clancy, D. J. 2008. Variation in mitochondrial genotype has substantial lifespan effects which may be modulated by nuclear background. Aging Cell 7:795–804.

Clancy, D. J., G. R. Hime, and A. D. Shirras. 2011. Cytoplasmic male sterility in Drosophila melanogaster associated with a mitochondrial CYTB variant. Heredity 107:374–376.

Clark, A. M. 1972. Influence of nutritional state of females on frequency of X/O males recovered after matings with irradiated males. Dros. Inf. Service 49:75.

Cooke, M. S., M. D. Evans, M. Dizdaroglu, and J. Lunec. 2003. Oxidative DNA damage: Mechanisms, mutation, and disease. FASEB J. 17:1195–1214.

Dean, R., B. Lemos, and D. K. Dowling. 2015. Context-dependent effects of Y chromosome and mitochondrial haplotype on male locomotive activity in Drosophila melanogaster. J. Evol. Biol. 28:1861–1871.

Denver, D. R., K. Morris, M. Lynch, L. L. Vassilieva, and W. K. Thomas. 2000. High direct estimate of the mutation rate in the mitochondrial genome of Caenorhabditis elegans. Science 289:2342–2344.

Dobler, R., B. Rogell, F. Budar, and D. K. Dowling. 2014. A meta-analysis of the strength and nature of cytoplasmic genetic effects. J. Evol. Biol. 27:2021–2034.

dos Santos, G., A. J. Schroeder, J. L. Goodman, V. B. Strelets, M. A. Crosby, J. Thurmond, D. B. Emmert, W. M. Gelbart, and t. F. Consortium. 2015. FlyBase: introduction of the Drosophila melanogaster Release 6 reference genome assembly and large-scale migration of genome annotations. Nucleic Acids Res. 43:D690–D697.

Dowling, D. K., K. C. Abiega, and G. Arnqvist. 2007a. Temperature-specific outcomes of cytoplasmic-nuclear interactions on egg-to-adult development time in seed beetles. Evolution 61:194–201.

Dowling, D. K., U. Friberg, F. Hailer, and G. Arnqvist. 2007b. Intergenomic epistasis for fitness: within-population interactions between cytoplasmic and nuclear genes in Drosophila melanogaster. Genetics 175:235–244.

Dowling, D. K., U. Friberg, and J. Lindell. 2008. Evolutionary implications of non-neutral mitochondrial genetic variation. Trends Ecol. Evol. 23:546–554.

Dowling, D. K., D. M. Tompkins, and N. J. Gemmell. 2015. The Trojan Female Technique for pest control: A candidate mitochondrial mutation confers low male fertility across diverse nuclear backgrounds in Drosophila melanogaster. Evol. Appl. 8:871–880.

Drake, J. W., B. Charlesworth, D. Charlesworth, and J. F. Crow. 1998. Rates of spontaneous mutation. Genetics 148:1667–1686.

Ellison, C. K. and R. S. Burton. 2006. Disruption of mitochondrial function in interpopulation hybrids of Tigriopus californicus. Evolution 60:1382–1391.

Frank, S. A. and L. D. Hurst. 1996. Mitochondria and male disease. Nature 383:224–225.

Gemmell, N. J., V. J. Metcalf, and F. W. Allendorf. 2004. Mother’s curse: The effect of mtDNA on individual fitness and population viability. Trends Ecol. Evol. 19:238–244.

Graf, U., M. M. Green, and F. E. Würgler. 1979. Mutagen-sensitive mutants in Drosophila melanogaster: Effects of premutational damage. Mutat. Res. Fund. Mol. Mech. Mut. 63:101–112.

Hadfield, J. D. 2010. MCMC methods for multi-response generalized linear mixed models: The MCMCglmm R package. J. Stat. Softw. 33:22.

Hill, G. E. 2015. Mitonuclear Ecology. Mol. Biol. Evol. 32:1917–1927.

Innocenti, P., E. H. Morrow, and D. K. Dowling. 2011. Experimental evidence supports a sex-specific selective sieve in mitochondrial genome evolution. Science 332:845–848.

Itsara, L. S., S. R. Kennedy, E. J. Fox, S. Yu, J. J. Hewitt, M. Sanchez-Contreras, F. Cardozo-Pelaez, and L. J. Pallanck. 2014. Oxidative stress is not a major contributor to somatic mitochondrial DNA mutations. PLoS Genet. 10:e1003974.

Kauppila, J. H. and J. B. Stewart. 2015. Mitochondrial DNA: Radically free of free-radical driven mutations. Acta Biochim. Biophys. Sin. 1847:1354–1361.

Keightley, P. D. and M. Lynch. 2003. Toward a realistic model of mutations affecting fitness. Evolution 57:683–689.

Larsson, N. G. 2010. Somatic mitochondrial DNA mutations in mammalian aging. Annu. Rev. Biochem. 79:683–706.

Latorre-Pellicer, A., R. Moreno-Loshuertos, A. V. Lechuga-Vieco, F. Sánchez-Cabo, C. Torroja, R. Acín-Pérez, E. Calvo, E. Aix, A. González-Guerra, A. Logan, M. L. Bernad-Miana, E. Romanos, R. Cruz, S. Cogliati, B. Sobrino, Á. Carracedo, A. Pérez-Martos, P. Fernández-Silva, J. Ruíz-Cabello, M. P. Murphy, I. Flores, J. Vázquez, and J. A. Enríquez. 2016. Mitochondrial and nuclear DNA matching shapes metabolism and healthy ageing. Nature 535:561–565.

Liu, H., Y. Jia, X. Sun, D. Tian, L. D. Hurst, and S. Yang. 2017. Direct determination of the mutation rate in the bumblebee reveals evidence for weak recombination-associated mutation and an approximate rate constancy in insects. Mol. Biol. Evol. 34:119–130.

Maklakov, A. A., U. Friberg, D. K. Dowling, and G. Arnqvist. 2006. Within-population variation in cytoplasmic genes affects female life span and aging in Drosophila melanogaster. Evolution 60:2081–2086.

Maklakov, A. A., S. Immler, H. Løvlie, I. Flis, and U. Friberg. 2013. The effect of sexual harassment on lethal mutation rate in female *Drosophila melanogaster*. Proc. R. Soc. Lond. B Biol. Sci. 280.

Mossman, J. A., L. M. Biancani, and D. M. Rand. 2016. Mitonuclear epistasis for development time and its modification by diet in *Drosophila*. Genetics 10.1534/genetics.116.187286.

Nabholz, B., S. Glemin, and N. Galtier. 2008. Strong variations of mitochondrial mutation rate across mammals‐‐the longevity hypothesis. Mol. Biol. Evol. 25:120–130.

Nabholz, B., S. Glémin, and N. Galtier. 2009. The erratic mitochondrial clock: variations of mutation rate, not population size, affect mtDNA diversity across birds and mammals. BMC Evol. Biol. 9:54.

Patel, M. R., G. K. Miriyala, A. J. Littleton, H. Yang, K. Trinh, J. M. Young, S. R. Kennedy, Y. M. Yamashita, L. J. Pallanck, and H. S. Malik. 2016. A mitochondrial DNA hypomorph of cytochrome oxidase specifically impairs male fertility in *Drosophila melanogaster*. eLife 5:e16923.

Pischedda, A. and A. Chippindale. 2005. Sex, mutation and fitness: asymmetric costs and routes to recovery through compensatory evolution. J. Evol. Biol. 18:1115–1122.

Rand, D. M., R. A. Haney, and A. J. Fry. 2004. Cytonuclear coevolution: The genomics of cooperation. Trends Ecol. Evol. 19:645–653.

Rogell, B., R. Dean, B. Lemos, and D. K. Dowling. 2014. Mito-nuclear interactions as drivers of gene movement on and off the X-chromosome. BMC Genomics 15:330.

Schrider, D. R., D. Houle, M. Lynch, and M. W. Hahn. 2013. Rates and genomic consequences of spontaneous mutational events in *Drosophila melanogaster*. Genetics 194:937–954.

Shadel, Gerald S. and Tamas L. Horvath. 2015. Mitochondrial ros signaling in organismal homeostasis. Cell 163:560–569.

Sharp, N. P. and A. F. Agrawal. 2008. Mating density and the strength of sexual selection against deleterious alleles in *Drosophila melanogaster*. Evolution 62:857–867.

Sharp, N. P. and A. F. Agrawal. 2012. Evidence for elevated mutation rates in low-quality genotypes. Proc. Natl. Acad. Sci. U. S. A. 109:6142–6146.

Sharp, N. P. and A. F. Agrawal. 2016. Low genetic quality alters key dimensions of the mutational spectrum. PLoS Biol. 14:e1002419.

Shaw, F. H. and C. F. Baer. 2011. Fitness-dependent mutation rates in finite populations. J. Evol. Biol. 24:1677–1684.

Smith, S., C. Turbill, and F. Suchentrunk. 2010. Introducing *Mother’s curse*: Low male fertility associated with an imported mtDNA haplotype in a captive colony of brown hares. Mol. Ecol. 19:36–43.

Sniegowski, P. D., P. J. Gerrish, T. Johnson, and A. Shaver. 2000. The evolution of mutation rates: separating causes from consequences. BioEssays 22:1057–1066.

Sollunn, S. J. and Ö. Strömnæs. 1964. The effect of temperature during irradiation on induced mutation frequency in *Drosophila melanogaster* sperm. Hereditas 51:10.1111/j.1601-5223.1964.tb01918.x.

Stewart, A. D., E. H. Morrow, and W. R. Rice. 2005. Assessing putative interlocus sexual conflict in *Drosophila melanogaster* using experimental evolution. Proc. R. Soc. Biol. Sci. Ser. B 272:2029–2035.

Vogel, E. W., R. L. Dusenbery, and P. D. Smith. 1985. The relationship between reaction kinetics and mutagenic action of monofunctional alkylating agents in higher eukaryotic systems: IV. The effects of the excision-defective mei-9L1 and mus(2)201D1 mutants on alkylation-induced genetic damage in *Drosophila*. Mutat. Res. Fund. Mol. Mech. Mut. 149:193–207.

Wang, A. D. and A. F. Agrawal. 2012. Dna repair pathway choice is influenced by the health of *Drosophila melanogaster*. Genetics 192:361–370.

Wolff, J. N., M. F. Camus, D. J. Clancy, and D. K. Dowling. 2015. Complete mitochondrial genome sequences of thirteen globally sourced strains of fruit fly (*Drosophila melanogaster*) form a powerful model for mitochondrial research. Mitochondrial DNA:4672–4674.

Wolff, J. N., E. D. Ladoukakis, J. A. Enriquez, and D. K. Dowling. 2014. Mitonuclear interactions: Evolutionary consequences over multiple biological scales. Philos. Trans. R. Soc. Lond. B Biol. Sci. 369:10.1098/rstb.2013.0443.

Wolff, J. N., N. Pichaud, M. F. Camus, G. Côté, P. U. Blier, and D. K. Dowling. 2016a. Evolutionary implications of mitochondrial genetic variation: Mitochondrial genetic effects on OXPHOS respiration and mitochondrial quantity change with age and sex in fruit flies. J. Evol. Biol. 29:736–747

Wolff, J. N., D. M. Tompkins, N. J. Gemmell, and D. K. Dowling. 2016b. Mitonuclear interactions, mtDNA-mediated thermal plasticity, and implications for the Trojan Female Technique for pest control. Sci. Rep. 6:30016.

Würgler, F. E. and P. Maier. 1972. Genetic control of mutation induction in *Drosophila melanogaster* I. Sex-chromosome loss in X-rayed mature sperm. Mutat. Res. Fund. Mol. Mech. Mut. 15:41–53.

Yee, W. K., K. L. Sutton, and D. K. Dowling. 2013. In vivo male fertility is affected by naturally occurring mitochondrial haplotypes. Curr. Biol. 23:R55–56.

Yee, W. K. W., B. Rogell, B. Lemos, and D. K. Dowling. 2015. Intergenomic interactions between mitochondrial and Y-linked genes shape male mating patterns and fertility in *Drosophila melanogaster*. Evolution 69:2876–2890

